# Prophage and metabolic determinants of *Staphylococcus aureus* survival to vancomycin identified via TraDIS screening

**DOI:** 10.64898/2026.05.25.727636

**Authors:** Sophia Zborowsky, Urszula Łapińska, Paul A. O’Neill, Audrey Farbos, Aaron R. Jeffries, Xiaoliang Ba, Mark A. Holmes, Maisem Laabei, Bing Zhang, Mark A. T. Blaskovich, Andrew Grant, Stefano Pagliara

**Affiliations:** Living Systems Institute, University of Exeter, Exeter, Devon, EX4 4QD, UK; Biosciences, University of Exeter, Exeter, Devon, EX4 4QD, UK; University of Exeter Sequencing Facility, University of Exeter, Exeter, Devon, EX1 2LU, UK; Department of Veterinary Medicine, University of Cambridge, Cambridge, CB3 0ES, UK; School of Cellular and Molecular Medicine, University of Bristol, Bristol, BS8 1TD, UK; Centre for Superbug Solutions, Institute for Molecular Bioscience, The University of Queensland, St. Lucia, QLD, 4072, Australia

## Abstract

Reduced vancomycin susceptibility phenotypes in *Staphylococcus aureus* contribute to treatment failure, yet the genetic determinants of survival under inhibitory vancomycin exposure remain incompletely defined. We performed transposon directed insertion-site sequencing (TraDIS) on a methicillin resistant *S. aureus* (MRSA) ST398 mutant library following exposure to vancomycin at its minimum inhibitory concentration, identifying 52 genes whose disruption was associated with loss of population survival at inhibitory drug concentrations. Prophage associated loci were the largest functional group, spanning predicted structural and regulatory genes as well as multiple conserved hypothetical proteins. Targeted testing of defined transposon mutants in a USA300 background confirmed that disruption of selected loci impaired growth under vancomycin exposure. Our results highlight the contribution of diverse physiological processes, including metabolism, stress responses, and a prominent role for prophage-associated functions, rather than discrete resistance pathways. Together, these findings indicate that vancomycin tolerance is shaped by the general physiological state of the bacterial cell, including metabolic capacity and stress adaptation.

**Importance:** Treatment failure in *Staphylococcus aureus* infections often occurs in the absence of known antibiotic resistance determinants, suggesting that additional survival mechanisms influence therapeutic outcomes. In this study, we identify genetic determinants required for survival during inhibitory vancomycin exposure, revealing a broad role for metabolic functions, stress adaptation, and prophage-associated loci. The prominence of these diverse processes highlights that survival reflects global physiological adaptation rather than discrete resistance pathways. This insight underscores the need to consider cellular physiology and stress responses when developing strategies to prevent antibiotic tolerance and improve treatment efficacy.

## Introduction

*Staphylococcus aureus* (*S. aureus*) is a major cause of invasive infections worldwide, ranging from skin and soft tissue infections to bacteremia, endocarditis, and pneumonia, and remains associated with substantial morbidity and mortality despite the availability of antibiotic therapy (1, 2). Treatment failure is frequently observed in *S. aureus* infections and cannot be fully explained by classical antibiotic resistance, suggesting that additional bacterial survival strategies contribute to poor therapeutic outcomes (3, 4).

One such strategy is antibiotic tolerance, defined as the ability of bacteria to survive prolonged exposure to otherwise lethal antibiotic concentrations through metabolic alterations (3, 5, 6). In *S. aureus*, tolerance and persistence phenotypes have been linked to a variety of physiological states, including altered metabolism, reduced growth rate, stress response activation, and biofilm formation, and have been shown to contribute to antibiotic treatment failure in both experimental models and clinical settings (1, 4, 7–11). These phenotypes are often transient and environment-dependent, complicating their detection and mechanistic dissection.

Vancomycin remains a key therapeutic option for severe infections caused by methicillin resistant *S. aureus* (MRSA). However, reduced susceptibility phenotypes such as heterogeneous vancomycin intermediate *S. aureus* (hVISA) and vancomycin intermediate *S. aureus* (VISA) have emerged and are associated with treatment failure (7, 12–14). VISA phenotypes typically arise through chromosomal mutations affecting global regulatory systems and cell envelope stress responses, leading to altered cell wall architecture, reduced autolysis, and impaired antibiotic penetration (12–14). Notably, many studies report correlations between altered gene expression levels and vancomycin susceptibility, while the underlying causal mutations and their functional consequences remain incompletely defined (8, 14).

Genome wide transposon insertion sequencing approaches, including TraDIS and Tn-seq, provide a powerful means to identify genes that contribute to bacterial fitness under defined selective pressures, including antibiotic stress (15, 16). By comparing transposon insertion patterns before and after exposure to antibiotics, these methods can identify genes that are dispensable for growth under normal growth conditions but become important for survival under antibiotic challenge. In *S. aureus*, and other bacteria, such approaches have been successfully used to investigate survival in host environments, stress adaptation, and antibiotic responses (17–22).

In this study, we used TraDIS to identify genes associated with survival of an *S. aureus* ST398 transposon library during exposure to vancomycin at the wild-type (WT) minimum inhibitory concentration (MIC). By focusing on survival at inhibitory concentrations rather than selection for resistant mutants, we aimed to identify determinants of antibiotic stress tolerance and fitness. Using genome wide transposon sequencing and targeted genetic perturbations, we identify multiple genes associated with *S. aureus* survival under inhibitory vancomycin exposure including a key envelope stress regulator, prophage and metabolism genes.

## Results and discussion

### Transposon library analysis identifies candidate genetic determinants for survival to vancomycin

The transposon mutant library used in this study was generated in *Staphylococcus aureus* sequence type 398 (ST398) strain S0385, a member of the LA-CC398 methicillin-resistant *S. aureus* (MRSA) lineage, a clonal complex originally identified in humans that has since become an expanding and concerning cause of zoonotic and human infections. (17). Although vancomycin remains a “last-resort” therapy for infections caused by CC398 clones, concerns persist that its widespread use, together with this lineage’s capacity to acquire resistance determinants, may drive reduced susceptibility, increasing the risk of treatment failure (23, 24). Exponential phase cultures of the library were exposed to vancomycin at the WT minimal inhibitory concentration (MIC; 1 µg/mL) or at 10× the MIC value, and population growth was monitored over time via optical density measurements. We used exponentially growing cultures because *S. aureus* exhibits intrinsic tolerance to vancomycin in stationary phase, where slow growth reduces antibiotic killing, making stationary-phase cultures less informative for identifying mutants affecting survival under drug stress (3, 25). When vancomycin was used at 10× the MIC value, no growth was observed for the transposon library in any replicate (Fig. 1A), indicating that vancomycin resistant mutants were not selected under these conditions. In contrast, when vancomycin was used at the MIC concentration, some cultures did not exhibit growth, whereas others exhibited limited growth across biological replicates (Fig. 1A). This growth was substantially reduced relative to drug free conditions, indicating that the transposon library exhibits limited population level proliferation during exposure to vancomycin. Consistent with this, time-kill assays (26) of exponentially growing bacteria (WT and transposon library) showed slow reduction in viable cell counts during vancomycin exposure, without evidence of neither regrowth over the assay period nor biphasic killing, symptom of persisters within susceptible populations (Fig. 1B). Together, these observations are consistent with the presence of vancomycin-intermediate or tolerant subpopulations within the mutant pool, rather than the emergence of fully resistant mutants.

**Figure 1.**
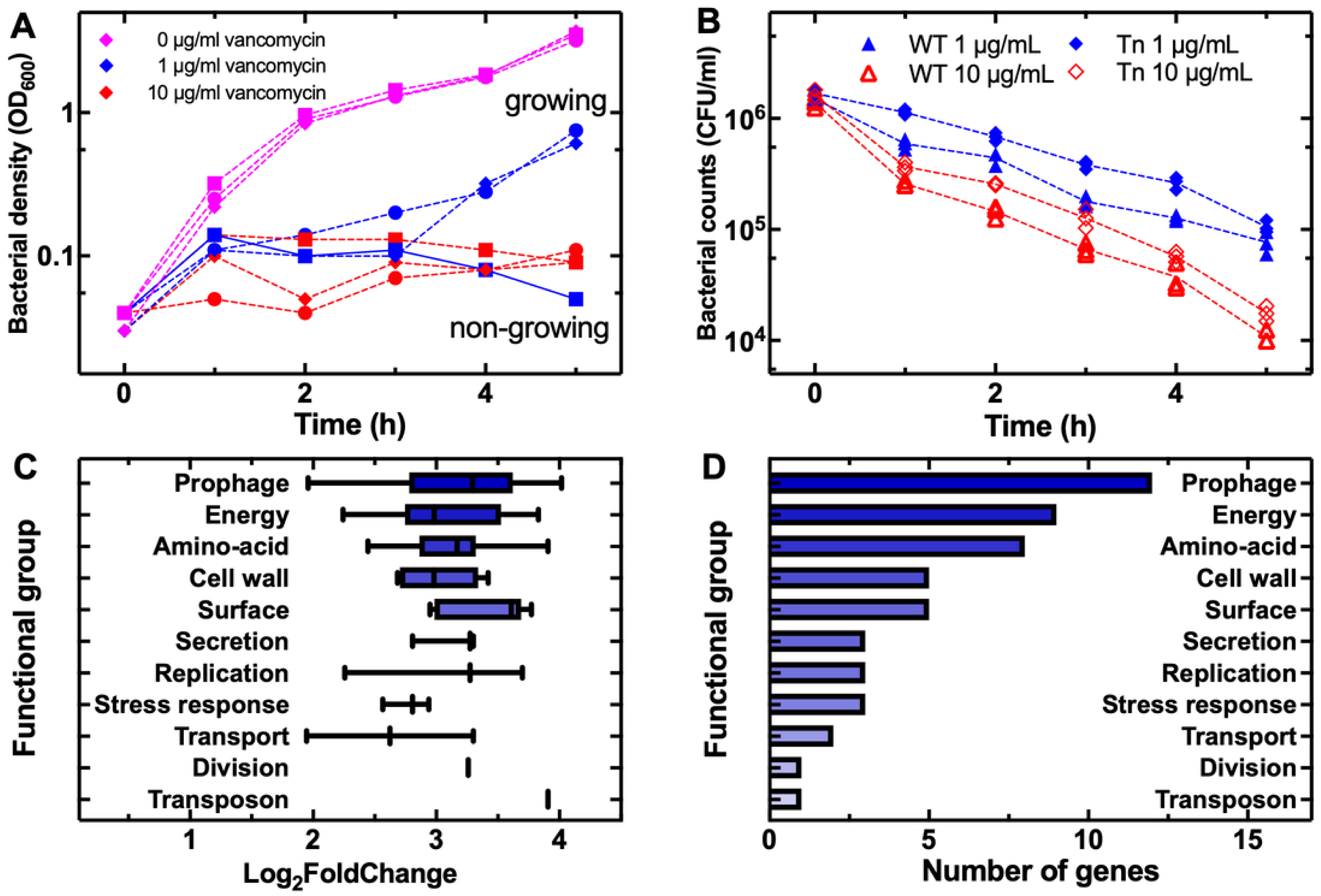
Growth of the *S. aureus* S0385 transposon library under vancomycin exposure and functional distribution of genes associated with reduced survival. (A) Growth curves of the S0385 transposon library measured by OD₆₀₀ in the absence of antibiotic (magenta data), at 1 µg/mL vancomycin (1× MIC, blue data), and at 10 µg/mL vancomycin (10× MIC, red data). Data represent three biological replicates, indicated by different symbols. **(B)** Time-kill assay for WT S0385 (triangles) and Tn S0385 (culture A, Table 1, diamonds) at 1 and 10 µg/mL vancomycin (blue and red data, respectively). Three biological replicas displayed. **(C)** Distribution of log₂ fold change values for genes within each functional category, shown as box-and-whisker plots. Boxes indicate the interquartile range, the central line denotes the median, and whiskers represent the minimum and maximum values. **(D)** Number of genes associated with loss of growth under vancomycin exposure, grouped by predicted biological function.

**Table 1.** Summary of transposon insertion coverage in TraDIS libraries following exposure to vancomycin at the MIC. statistics detailing transposon insertion coverage across TraDIS mutant libraries derived from cultures exposed to vancomycin at the MIC. Libraries A-C represent populations that grew under vancomycin exposure, while libraries D-F represent populations that did not grow under the same conditions. For each library, the total number of transposon insertions detected, the number of genes containing at least four insertions, the percentage of genes with insertions relative to the total number of genes in the genome (n = 2,498), and the average number of insertions per gene are provided.

| Culture | Total unique insertions | Genes with $\geq 4$ insertions | % genes with $\geq 4$ insertions | Mean insertions per gene |
| --- | --- | --- | --- | --- |
| **A – Growth** | 161,810 | 2,431 | 97.32% | 64.8 |
| **B – Growth** | 88,996 | 2,315 | 92.67% | 35.6 |
| **C – Growth** | 6,385 | 621 | 24.86% | 2.6 |
| **D – No growth** | 246,115 | 2,492 | 99.76% | 98.5 |
| **E – No growth** | 215,568 | 2,488 | 99.60% | 86.3 |
| **F – No growth** | 274,540 | 2,497 | 99.96% | 90.0 |

Based on these outcomes, cultures that exhibited detectable growth at the vancomycin MIC were designated “growing”, whereas cultures that failed to grow under the same conditions were designated “non-growing” (Fig. 1A). The differential growth outcomes across biological replicates likely reflect stochastic sampling effects inherent to transposon libraries, where each experiment captures only a fraction of the total mutant diversity; consequently, some cultures may include mutants capable of surviving at the MIC, whereas others may not.

To identify genes associated with survival during vancomycin exposure at its MIC, cultures of the transposon library containing 10⁹ cells were exposed to vancomycin (1 µg/mL) for five hours. This inoculum size was selected to ensure that at least 10⁸ cells remained at the end of the experiment, providing greater than 100× coverage of the ∼250,000 unique insertion sites present in the library (17). This exceeds the ∼2.5 × 10⁷ cells required for 100× coverage and ensures sufficient representation of mutants, as well as adequate genomic DNA yield for downstream library preparation and sequencing. Genomic DNA was extracted from three growing and three non-growing cultures and subjected to transposon-directed insertion-site sequencing (TraDIS) using a transposon-specific primer (see methods) to map insertion sites genome-wide. Sequencing reads were mapped to the reference genome of *S. aureus* strain S0385 (GeneBank AM990992.1), insertion counts were quantified for each gene, and differential insertion analysis was performed between the growing and non-growing cultures (File S1).

Genes disrupted with transposons were considered significant if they exhibited a log₂ fold change (log_2_FC) greater than 1.5 and a statistically significant p value and were not essential for growth in LB alone. We note that genes classified as essential for growth in LB (File S2) may also contribute to vancomycin tolerance; however, because insertions in these genes are depleted in both growing and non-growing cultures, their contribution could not be distinguished in this comparative framework, and they were therefore excluded from downstream analyses.

TraDIS sequencing of growing cultures identified between 6,385 and 274,540 transposon insertions per replicate (Table 1). Insertions were detected in 92–99% of genes across five independent culture replicates derived from the master transposon library, while one of the six cultures showed lower coverage (Table 1). Overall, these data indicate near-complete genome coverage across the majority of replicates. Cultures propagated under vancomycin exposure had fewer insertions than unselected controls, likely due to a combination of population bottlenecks and selective pressure.

This analysis identified 52 genes that were significantly depleted in non-growing cultures relative to growing cultures (Table 2 and File S3). Disruption of these genes was therefore associated with loss of proliferation under vancomycin exposure. No genes were found to be significantly enriched in non-growing cultures, indicating that we detected no evidence for genes whose disruption reproducibly enhanced growth under vancomycin exposure in this experimental setting.

**Table 2.**
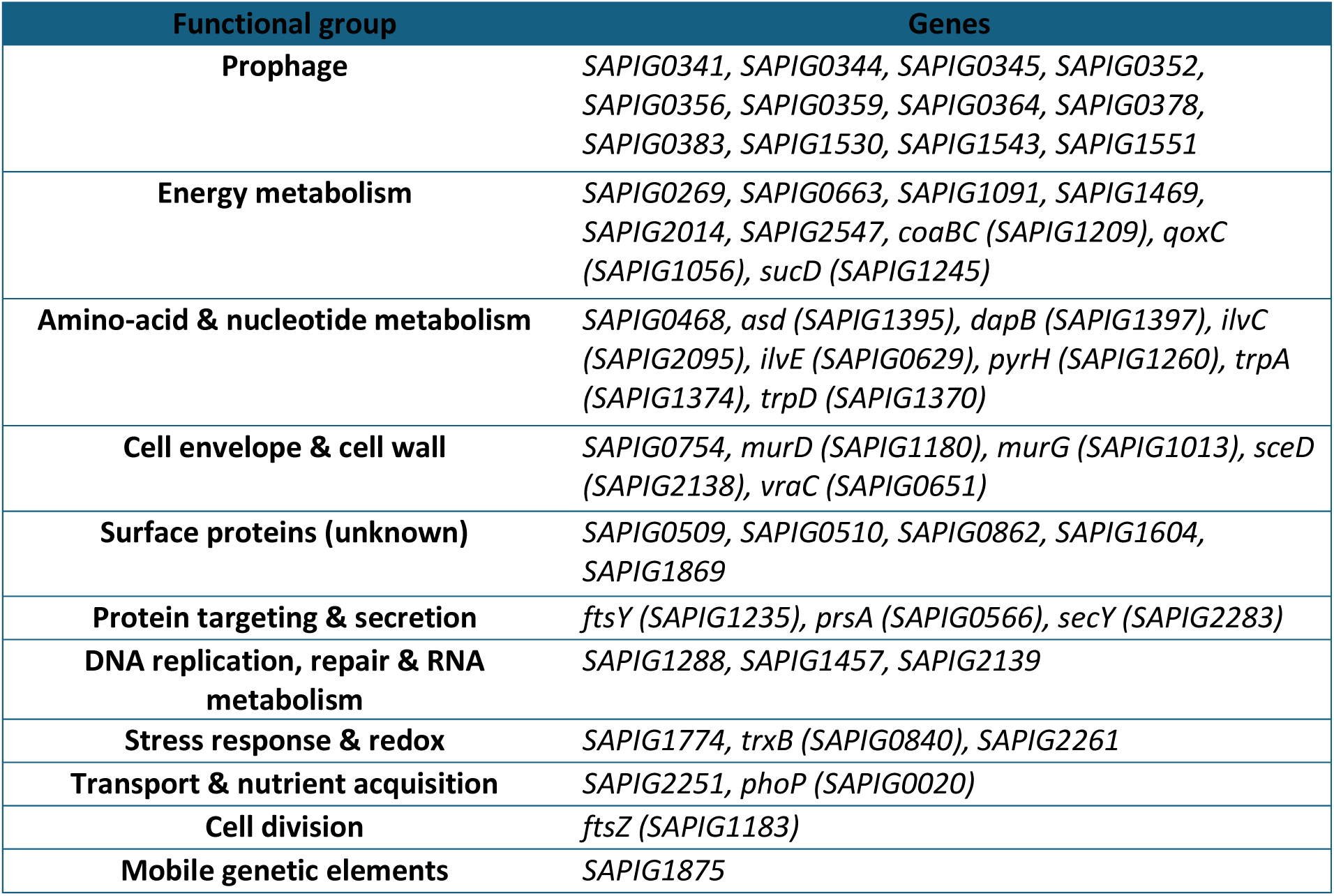
Functional grouping of genes associated with loss of survival at the vancomycin MIC. Genes identified by TraDIS as significantly enriched in MIC-non-surviving cultures are grouped by predicted biological function based on genome annotation. Gene names are shown where available; otherwise, locus tags are provided. Genes were considered significant based on the following criteria: log₂FC > 1.5, a statistically significant p value, ≤3 transposon insertions in genes required for survival at the vancomycin MIC (File S1), and ≥4 insertions in at least one replicate under LB growth conditions, indicating that the gene is not required for growth in LB (File S2).

| Functional group | Genes |
| --- | --- |
| **Prophage** | *SAPIG0341, SAPIG0344, SAPIG0345, SAPIG0352, SAPIG0356, SAPIG0359, SAPIG0364, SAPIG0378, SAPIG0383, SAPIG1530, SAPIG1543, SAPIG1551* |
| **Energy metabolism** | *SAPIG0269, SAPIG0663, SAPIG1091, SAPIG1469, SAPIG2014, SAPIG2547, coaBC (SAPIG1209), qoxC (SAPIG1056), sucD (SAPIG1245)* |
| **Amino-acid & nucleotide metabolism** | *SAPIG0468, asd (SAPIG1395), dapB (SAPIG1397), ilvC (SAPIG2095), ilvE (SAPIG0629), pyrH (SAPIG1260), trpA (SAPIG1374), trpD (SAPIG1370)* |
| **Cell envelope & cell wall** | *SAPIG0754, murD (SAPIG1180), murG (SAPIG1013), sceD (SAPIG2138), vraC (SAPIG0651)* |
| **Surface proteins (unknown)** | *SAPIG0509, SAPIG0510, SAPIG0862, SAPIG1604, SAPIG1869* |
| **Protein targeting & secretion** | *ftsY (SAPIG1235), prsA (SAPIG0566), secY (SAPIG2283)* |
| **DNA replication, repair & RNA metabolism** | *SAPIG1288, SAPIG1457, SAPIG2139* |
| **Stress response & redox** | *SAPIG1774, trxB (SAPIG0840), SAPIG2261* |
| **Transport & nutrient acquisition** | *SAPIG2251, phoP (SAPIG0020)* |
| **Cell division** | *ftsZ (SAPIG1183)* |
| **Mobile genetic elements** | *SAPIG1875* |

The 52 genes associated with growth at the vancomycin MIC were assigned the protein they code and classified according to their predicted biological functions and cellular processes (Table 2, File S3). These genes all displayed a strong log_2_FC ranging between 1.9 and 4.0 (Fig. 1C). In the following sections, we describe these functional categories and discuss individual genes and pathways that may contribute to growth during exposure to vancomycin.

### Prophage associated genes are enriched among determinants of growth during treatment with vancomycin

The largest functional category identified in the TraDIS analysis comprised prophage associated genes, accounting for 12 of the 52 genes linked to growth during vancomycin treatment (Fig. 1D, Table 2, File S3). Prophage regions in the S0385 genome were identified using PHASTER (PHAge Search Tool Enhanced Release) based on the complete chromosomal sequence (GenBank accession AM990992.1), which predicted three distinct prophage regions.

Of the prophage associated genes identified in our TraDIS analysis, we found genes belonging to two of the three predicted phage regions only; nine genes mapped to a single predicted prophage region (locus tags SAPIG0341–SAPIG0383), while three mapped to a second predicted prophage region (File S3). The group of nine genes was annotated as homologous to genes encoded by *Staphylococcus* phage StauST398-2 (27, 28) (GenBank RefSeq NC_021323.1), a temperate phage originally described in ST398 backgrounds. In general, *S. aureus* strain S0385 harbors multiple prophage regions as part of its accessory genome, and variation in prophage content contributes to genomic differences among ST398 isolates (27, 28). Moreover, StauST398-2 related prophages have been associated with differences in virulence traits among ST398 isolates (29, 30). The homology of these genes to StauST398-2 therefore indicates close relatedness between the predicted prophage regions present in strain S0385 and previously characterized ST398 associated phages. The second predicted prophage region did not display homology to any known prophage.

The TraDIS identified prophage associated genes include predicted structural proteins, regulatory elements such as the integrase regulator RinB and a helix-turn-helix DNA binding protein, a prophage encoded dUTPase, and multiple hypothetical proteins (File S3). Pfam motif analysis further indicated that several of these proteins contain conserved phage related domains consistent with roles in virion assembly and phage regulation, including a capsid scaffolding protein, a head completion protein, and a tail assembly chaperone (File S4). Together, these annotations suggest that the identified loci represent components of functional prophage modules. However, many of the proteins remain annotated as conserved hypothetical proteins, and their specific roles in host physiology or antibiotic stress responses remain unknown.

We examined whether prophage associated genes identified in the TraDIS screen were transcriptionally upregulated during exposure to vancomycin. For this, WT *S. aureus* S0385 cultures were exposed to vancomycin at its MIC value, and cDNA levels of two predicted prophage genes from each of the two TraDIS identified regions (SAPIG059 and SAPIG378 for the StauST398-2-like prophage and SAPIG1530 and SAPIG1551 for the second predicted prophage) were quantified by RT-qPCR. We found that three out of four tested prophage associated genes were significantly upregulated upon vancomycin exposure compared to untreated controls (Fig. 2A).

**Figure 2.**
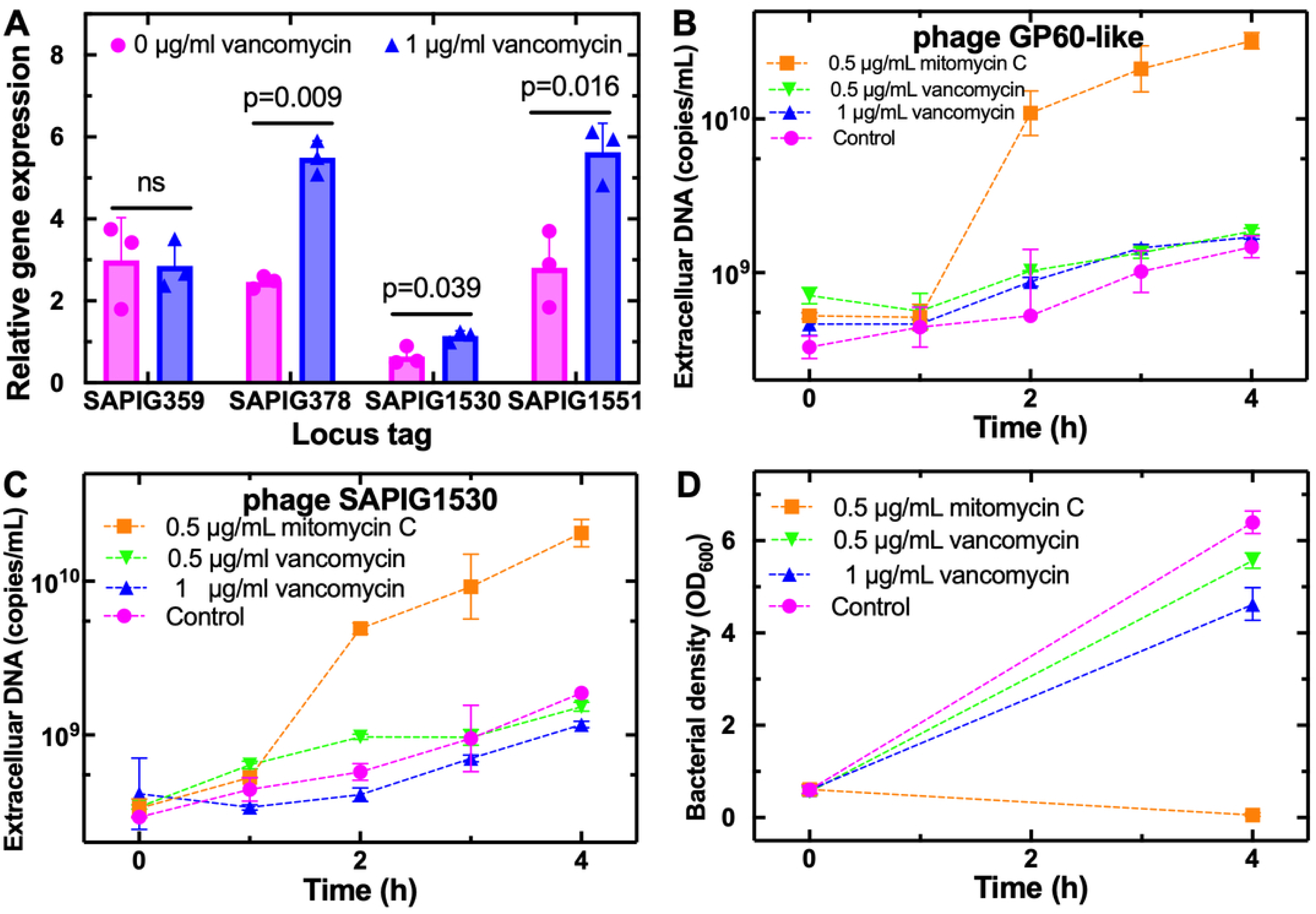
Prophage induction during exposure to vancomycin. **(A)** Expression of prophage associated genes in WT S0385 in the absence (magenta circles) and presence of 1 µg/ml vancomycin (blue triangles). Expression was measured by RT-qPCR and normalized to the gmk gene (n = 3). p values determined by Welch’s t-test are indicated in the graph. Points represent individual biological replicates and bars indicate mean and standard deviation. **(B)** Extracellular DNA for the predicted prophage region one quantified via qPCR by targeting a gp60-like phage gene following exposure to LB medium (magenta circles), treatment with mitomycin C (0.5 µg/mL, orange squares) or vancomycin (0.5 or 1 µg/mL, green downward triangles and blue upward triangles, respectively). The gp60-like gene was identified based on sequence homology to the gp60 gene of Staphylococcus phage StauST398-2 (NCBI Reference Sequence: NC_021323.1), a structural phage gene, and was used as a proxy for extracellular prophage DNA. **(C)** Extracellular DNA for the second predicted prophage region, using primers for the gene SAPIG1530 following exposure to LB, mitomycin C or vancomycin. **(D)** Bacterial density measured via OD₆₀₀ at 0 and 4 h for control cultures and cultures exposed to either mitomycin C or vancomycin. Data represent mean and standard deviation from three biological replicates at each time point.

We next examined prophage induction and release of viral particles from the StauST398-2-like prophage region and the second prophage predicted region in strain S0385 under antibiotic stress. We quantified extracellular phage gene copies while monitoring culture density (OD₆₀₀) following treatment with vancomycin or mitomycin C as a control. The StauST398-2-like prophage and the second predicted prophage were robustly induced by mitomycin C, as evidenced by a marked increase in extracellular phage DNA (Fig. 2B and 2C) and a decline in culture density (Fig. 2D), confirming that the prophages are functional and capable of entering the lytic cycle. In contrast, no increase in extracellular phage DNA or culture collapse, as indicated by density, was observed following vancomycin treatment (Fig. 2B, 2C and 2D), indicating that vancomycin exposure does not trigger lytic activation of this prophage under the conditions tested.

These results indicate that vancomycin exposure induces transcriptional activation of prophage associated genes. This is consistent with antibiotic mediated stress triggering of prophage gene expression under similar conditions in other strains (31, 32). Furthermore, vancomycin did not induce phage entry into the lytic cycle, suggesting that prophage activation under these conditions is limited to gene expression rather than productive phage replication. The induction of prophage encoded genes may therefore contribute to bacterial proliferation under vancomycin stress.

*S. aureus* prophages are recognized as major contributors to *S. aureus* genetic diversity, host adaptation, virulence, and resistance to antibiotics (28, 31–34). Furthermore, the vast majority of clinical isolates harbor one or more prophages, which encode a substantial fraction of the accessory genome and vary systematically across clinical contexts and host environments (28, 35). Despite these associations, mechanistic understanding of how prophage encoded genes contribute specifically to antibiotic tolerance, persistence, or survival remains limited, as most implicated prophage loci are poorly characterized and supported primarily by correlative genomic data (35). Our TraDIS analysis implicates multiple prophage-associated genes in growth under vancomycin exposure, yet the predominance of uncharacterized prophage genes among these “hits” underscores the current lack of mechanistic insight into how prophage regions influence antibiotic stress responses and emphasizes the need for further experimental investigation.

### Energy metabolism and nutrient acquisition genes contribute to growth in the presence of vancomycin

Genes involved in energy metabolism constituted the second largest functional category identified in the TraDIS analysis, accounting for 9 of the 52 genes associated with *S. aureus* growth at the vancomycin MIC (Fig. 1D, Table 2, File S3). These genes span distinct aspects of cellular energy generation and metabolic regulation.

To further assess the contribution of specific genes to vancomycin susceptibility, we used the Nebraska Transposon Mutant Library (NTML), a sequence defined transposon insertion collection generated in the *S. aureus* USA300 background (36) (File S5). Mutants were tested alongside the USA300 parental strain JE2 for vancomycin susceptibility using broth microdilution assays (37).

Vancomycin MIC values for the tested USA300 transposon mutants ranged from 1 to 2 µg/mL and were comparable to that of the parental JE2 strain (2 µg/mL; File S6). To assess vancomycin susceptibility beyond MIC measurements, growth kinetics were monitored at 1 µg/mL vancomycin (Fig. 3). All mutants grew comparably to the WT in the absence of antibiotics, indicating no baseline growth defect (Fig. S1A). At 2 µg/mL, no growth was observed for any strain (Fig. S1B).

**Figure 3.**
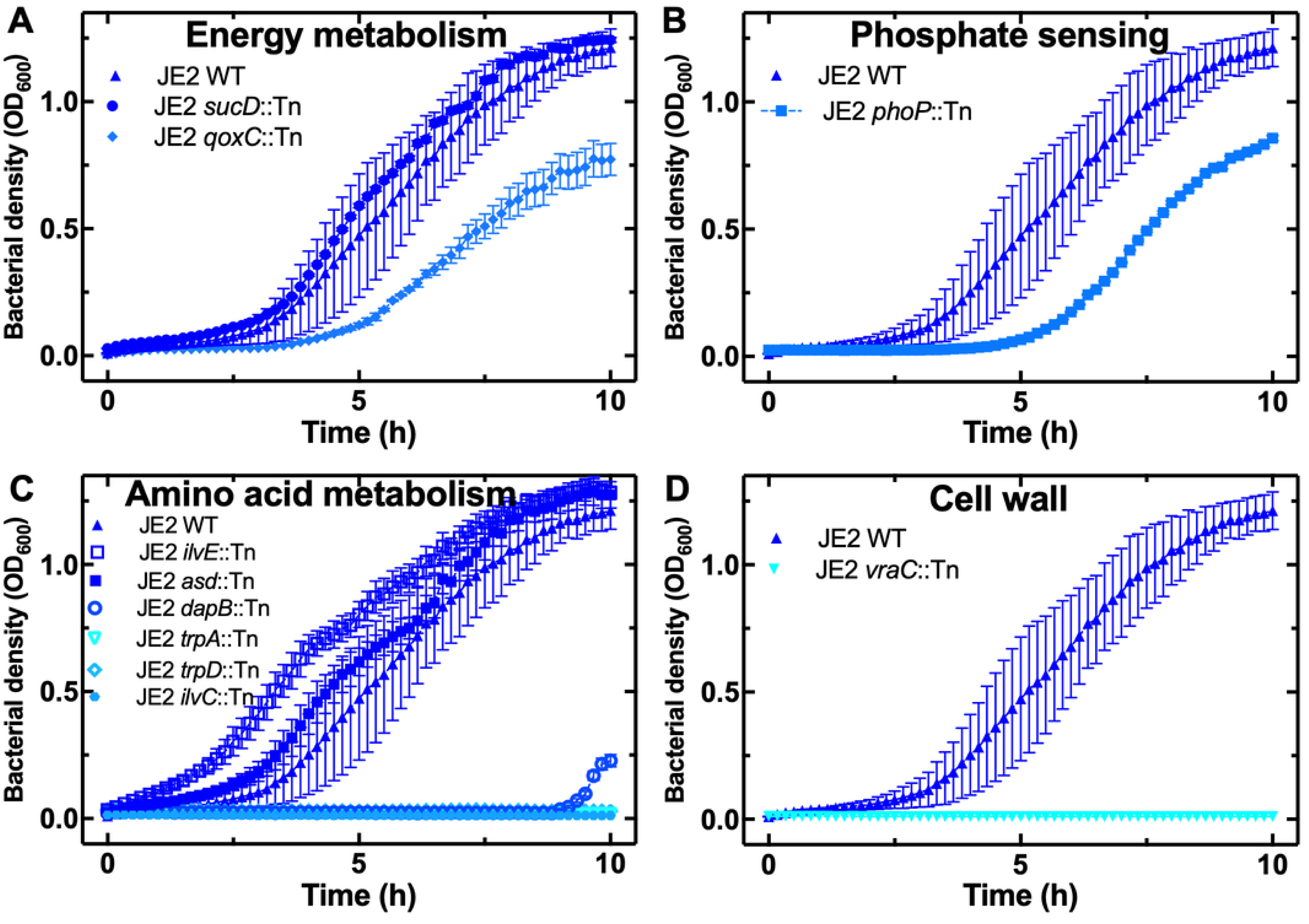
Impact of selected USA300 transposon mutants on growth during vancomycin exposure. Growth of defined mutants from the *S. aureus* USA300 JE2 Nebraska Transposon Mutant Library disrupted in genes involved in **(A)** energy metabolism, **(B)** phosphate sensing, **(C)** amino acid metabolism and **(D)** cell envelope stress response. Growth was monitored by OD₆₀₀ during exposure to vancomycin at a concentration of 1 µg/mL. Data represent mean and standard deviation from three biological replicates.

Disruption of *qoxC*, encoding a subunit of the cytochrome aa₃ terminal oxidase, resulted in impaired growth at 1 µg/mL vancomycin (i.e. 0.5 × the MIC value for the parental JE2 strain) compared to the WT strain, despite comparable growth in drug-free conditions (Fig. 3A, Fig. S1A). In contrast, disruption of *sucD*, encoding succinyl-CoA synthetase of the tricarboxylic acid cycle, did not alter growth under vancomycin exposure and behaved similarly to the WT across all tested concentrations (Fig. 3A, Fig. S1A). The vancomycin-dependent growth defect observed in the *qoxC* mutant suggests that respiratory capacity may contribute to bacterial fitness during cell wall inhibition, consistent with evidence that energy generation can influence antibiotic stress responses (38). By contrast, the absence of a detectable phenotype for the *sucD* mutant indicates that not all genes identified in the TraDIS screen confer measurable susceptibility phenotypes in the USA300 transposon library.

The TraDIS analysis also identified genes involved in nutrient uptake and phosphate regulation that were associated with survival at the vancomycin MIC. These included the global phosphate responsive regulator *phoP*. A *phoP* USA300 transposon mutant showed reduced growth during exposure to 1 µg/mL vancomycin (Fig. 3B). PhoP is a global regulatory protein that links phosphate availability to broad metabolic functions, including cell wall composition and virulence in *S. aureus* (39, 40). Disruption of *phoP* may therefore impair these protective mechanisms and increase antibiotic sensitivity; in this context, it is plausible that *phoP* knockout could contribute to enhanced susceptibility to vancomycin.

Furthermore, previous research has linked *phoP* to antibiotic resistance, *e.g*. in *Pseudomonas aeruginosa* the PhoP/PhoQ system plays a role in resistance to polymyxin B, colistin and antimicrobial peptides through the regulation of genes involved in cell envelope remodeling and lipid modification (41, 42).

### Amino acid biosynthesis and central metabolic pathways are associated with growth during exposure to vancomycin

A third prominent functional category identified in the TraDIS analysis comprised genes involved in amino acid biosynthesis and central metabolic pathways (8 out of 52 genes) (Fig. 1D, Table 2, File S3). This group included enzymes from multiple biosynthetic routes, including the aspartate family pathway (*asd*), lysine biosynthesis via the diaminopimelate (DAP) pathway (*dapB*), branched chain amino acid biosynthesis (*ilvC*, *ilvE*), and tryptophan biosynthesis (*trpA*, *trpD*).

Disruption of *asd, dapB, ilvE,* and *trpD* did not alter the MIC, while *ilvC* and *trpA* mutants exhibited a modest reduction in MIC (1 µg/mL) (File S6). Under exposure to 1 µg/mL vancomycin, the *asd* and *ilvE* mutants grew similarly to the WT, whereas *dapB, trpA, trpD,* and *ilvC* mutants failed to grow (Fig. 3C), indicating a vancomycin dependent fitness defect.

Disruption of a defined subset of amino acid biosynthesis genes produced consistent vancomycin susceptibility phenotypes across two *S. aureus* strains, supporting a link between these pathways and growth during exposure to vancomycin. Notably, multiple genes within the same biosynthetic pathways, particularly the tryptophan biosynthesis pathway (*trpA* and *trpD*), were identified both in the S0385 TraDIS screen and in USA300 transposon knockouts, strengthening the interpretation that this pathway is related to sustained growth in the presence of vancomycin.

Tryptophan biosynthesis and amino acid metabolism have previously been linked to bacterial stress responses and survival under antibiotic pressure through indirect effects on metabolic balance and global stress adaptation, rather than through direct interaction with antibiotic targets (43–45). On this basis, we speculate that disruption of these biosynthetic pathways limits the availability of specific amino acids required to sustain adaptive physiological responses during vancomycin induced stress in *S. aureus*, although the precise mechanisms remain to be determined.

### Cell envelope and cell wall metabolism genes are linked to growth during exposure to vancomycin

The TraDIS analysis identified a subset of genes involved in cell envelope stress responses and cell wall biogenesis that were associated with growth under exposure to vancomycin at its MIC value (Fig. 1D, Table 2, File S3). These genes included the cell envelope stress response regulator *vraC*, a putative acetyltransferase predicted to mediate cell wall modification (*SAPIG0754*), the cell wall remodeling enzyme *sceD*, and two peptidoglycan biosynthesis genes, *murG* and *murD*. For genes that are normally required for growth, such as *murG* and *murD* (46), transposon insertions detected in the TraDIS library likely represent insertions in regions that do not result in complete loss of function, permitting survival while still affecting fitness under vancomycin stress.

Additional phenotypic characterization was limited to *vraC* using a defined USA300 transposon mutant (File S5), given the lack of available defined mutants for the remaining loci.

Disruption of *vraC* in the USA300 background resulted in a reduced vancomycin MIC compared to the WT strain (File S6) and complete growth inhibition at 1 µg/mL vancomycin (Fig. 3D), indicating increased susceptibility to vancomycin upon loss of *vraC* function. This phenotype aligns with a previously reported *vraC* knockout in *S. aureus* strain Mu50, where it led to increased vancomycin susceptibility, accompanied by a reduction in cell-wall thickness (36).

We tested if knock out of *vraC* influences vancomycin binding by measuring the fluorescence of a fluorescent vancomycin derivative in individual *S. aureus* cells by flow cytometry (47). Exponentially growing cultures of *S. aureus* JE2 WT and the USA300 *vraC* knockout strain were incubated with a fluorescent derivative of vancomycin (*i.e*. vancomycin linked to the small fluorophore, nitrobenzoxadiazole -NBD- via click chemistry) (48). Aliquots were analyzed after 10 min, 1 h, or 4 h incubation. At the 10 min time point, both the WT and the USA300 *vraC* knockout populations rapidly accumulated vancomycin-NBD, with approximately 99% of cells scoring as fluorescence positive, indicating rapid binding, the *vraC* knockout population displaying slightly higher fluorescence. We repeated this test with all USA300 transposon mutants above and found similar rapid binding of labeled vancomycin, with the *trpA* knockout population displaying slightly higher fluorescence compared to the other strains (Fig. S2).

### Additional functional groups associated with survival under vancomycin exposure

In addition to the major functional categories identified above, several genes associated with growth during vancomycin exposure were identified for which no defined USA300 transposon mutants were available. These included predicted cell surface and envelope associated proteins of unknown function (SAPIG0509, SAPIG0510, SAPIG0862, SAPIG1869, and SAPIG1604), genes involved in protein targeting and secretion (*prsA, secY,* and *ftsY*), stress and redox regulation (SAPIG2261, *osmC/ohr* family protein, *trxB*), and a small number of genes linked to DNA replication, repair, and RNA metabolism (SAPIG2139, SAPIG1288, and SAPIG1457). Although these genes span diverse biological processes and lack direct functional validation in this study, their enrichment in the cultures growing in the presence of vancomycin suggests that maintenance of envelope integrity, protein handling, redox balance, and core information processing functions may contribute to bacterial fitness during vancomycin induced stress. Given the role of vancomycin in perturbing cell wall synthesis, disruption of these supporting systems may indirectly compromise the ability of cells to withstand antibiotic challenge. However, in the absence of additional phenotypic characterization, the specific contributions of these loci remain unresolved.

### Disruption of selected loci alters survival kinetics during vancomycin exposure

We performed time-kill assays to determine whether disruption of selected genes affects the tolerance phenotype, which was observed for the WT S0385 and the TraDIS library (Fig. 1B). The genes analyzed (*trpD*, *phoP*, *vraC*, *ilvC*, and *qoxC*) were selected because their disruption resulted in reduced growth under sub-inhibitory vancomycin conditions in the growth kinetics assay (Fig. 3), and they represent the different biological functional groups identified above. Exponentially growing cultures of the parental JE2 WT strain and selected mutants were exposed to vancomycin at the WT MIC (2 µg/mL), and viable cell counts were quantified over time.

The JE2 WT strain exhibited a gradual reduction in viable cell counts over the course of the assay, indicating slow killing kinetics under vancomycin exposure, similarly to WT S0385 (Fig. 4). No evidence of regrowth or biphasic killing was observed, consistent with a tolerance-like survival phenotype under these conditions. All selected mutants displayed increased susceptibility to vancomycin, relative to the WT (Fig. 4). These results strengthen the observations from Fig. 3, showing that genes whose disruption reduced growth in the presence of vancomycin also contribute to decreased survival under bactericidal vancomycin concentrations.

**Figure 4.**
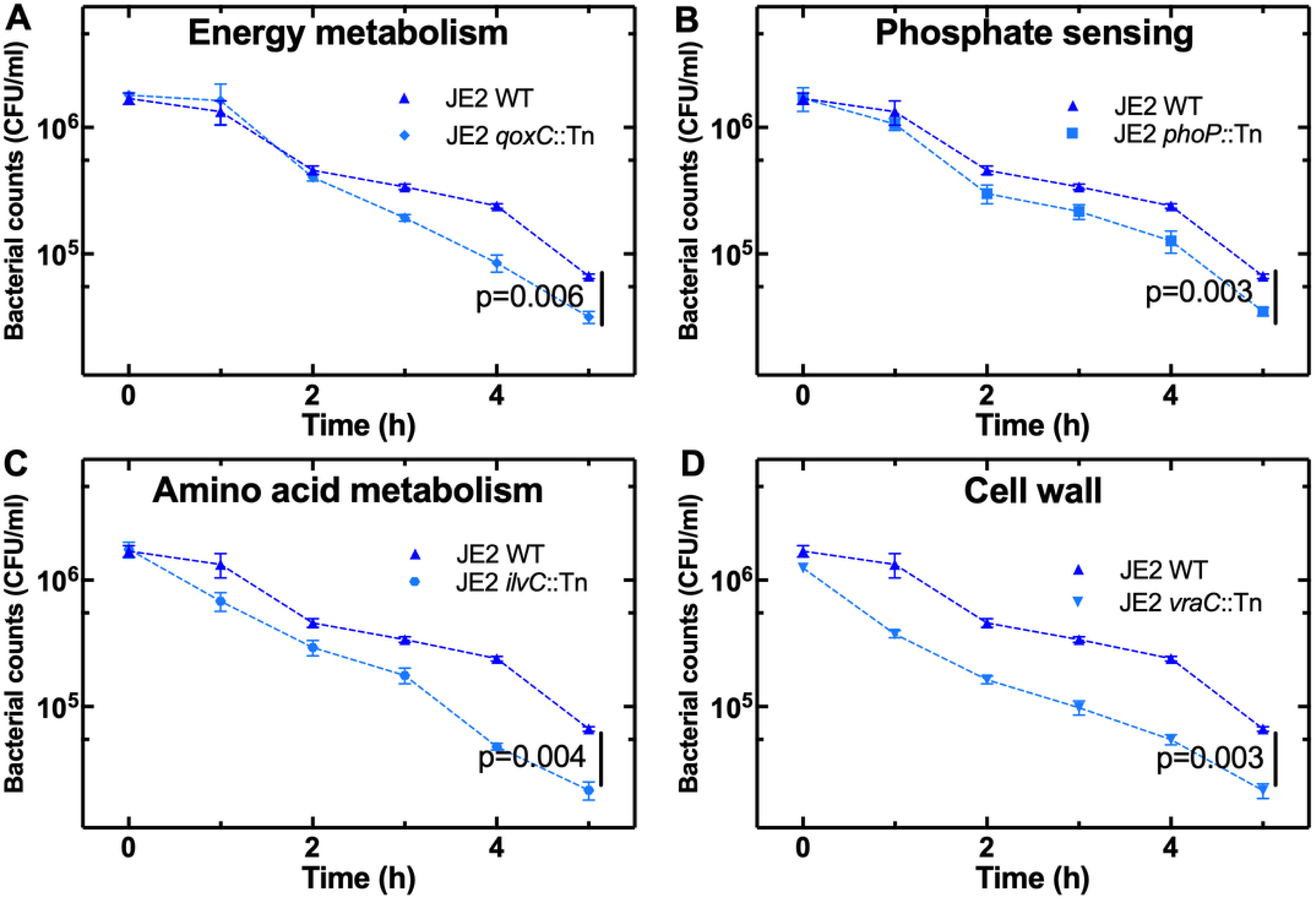
Impact of selected USA300 transposon mutants on survival during vancomycin exposure. Time-kill assays of *S. aureus* USA300 JE2 and selected transposon mutants. Exponentially growing cultures were exposed to vancomycin at 2 µg/mL (WT MIC), and viable cell counts (CFU/mL) were determined at the indicated time points. Strains shown include the parental JE2 strain and mutants disrupted in in **(A)** energy metabolism, **(B)** phosphate sensing, **(C)** amino acid metabolism and **(D)** cell envelope stress response. Data represent mean and standard deviation from three biological replicates. At t = 5 h, statistical significance was assessed using Welch’s t-test comparing WT (Group A) to each mutant.

### Cross-antibiotic growth assessment of vancomycin-selected TraDIS mutants

To determine whether the genes identified by the TraDIS sequencing as related to growth under inhibitory vancomycin conditions, were also involved in growth under other antibiotics, USA300 transposon mutants displaying reduced growth in the presence of vancomycin were exposed to sub-MIC (0.5× MIC) concentrations of antibiotics from different classes: penicillin G (β-lactam), ciprofloxacin (fluoroquinolone), and roxithromycin (macrolide). Exponentially growing cultures of the WT JE2 strain and selected USA300 transposon mutants were monitored for growth over time. All mutants exhibited reduced growth compared to the WT JE2 strain under all three antibiotics; specifically, the *dapB*, *ilvC* and *trpA* knockout mutants displayed the lowest growth (Fig. 5), further corroborating that amino acid biosynthesis is linked to survival under antibiotic pressure without direct interaction with specific antibiotic targets (43–45). In contrast, the *vraC* mutant was most affected by vancomycin and penicillin G corroborating the notion that this gene plays an important role in survival to cell wall targeting antibiotics (36).

**Figure 5.**
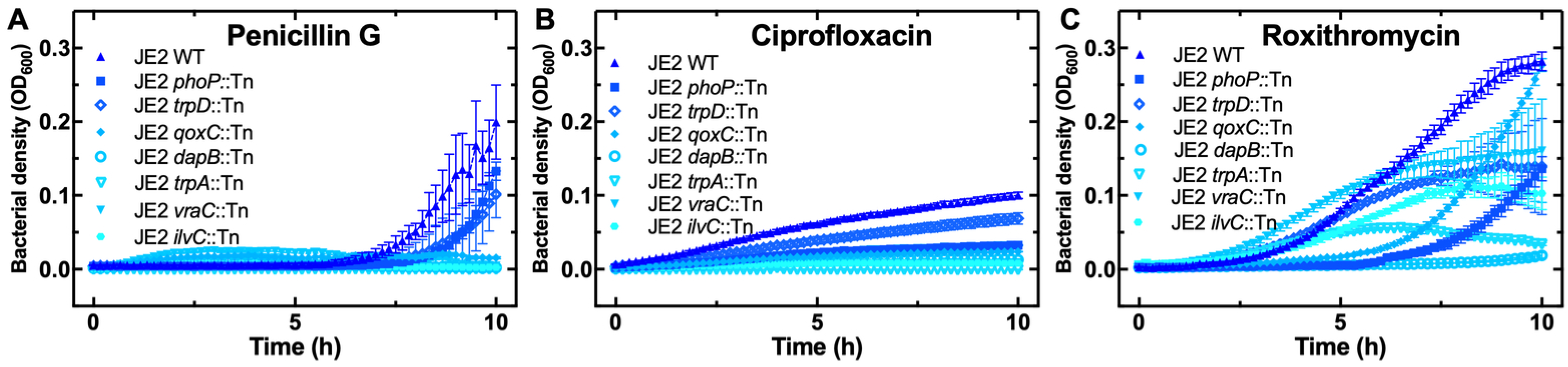
Impact of selected USA300 transposon mutants on growth during exposure to other antibiotics. Exponentially growing cultures of the WT JE2 strain and selected transposon mutants were exposed to sub-MIC concentrations of **(A)** penicillin G (2 µg/mL; 0.5× MIC of 4 µg/mL), **(B)** ciprofloxacin (32 µg/mL; 0.5× MIC of 64 µg/mL), and **(C)** roxithromycin (50 µg/mL; near 0.5× MIC, MIC >64 µg/mL). Growth was monitored over time by OD600. Data represent the mean and standard deviation of three biological replicates.

These results indicate that many of the genes important for growth under vancomycin stress also contribute to fitness under other antibiotic stresses. The further reduction in growth observed for the mutants suggests that these loci play broader roles in maintaining cellular functions required for survival during diverse antibiotic challenges. Consistent with previous studies, ciprofloxacin tolerance in *S. aureus* has been linked to global physiological adaptations involving metabolic reprogramming, including altered amino acid biosynthesis and energy metabolism (49). This supports the idea that disruption of genes affecting metabolism, envelope stress, or energy production, such as those identified in our TraDIS screen, can sensitize bacteria to multiple bactericidal antibiotics.

## Conclusions

In this study, we combined population-level TraDIS and individual defined transposon mutants to identify genetic determinants of *S. aureus* growth in the presence of vancomycin. By focusing on outcomes near the vancomycin MIC, we uncovered genes whose disruption compromises survival during antibiotic exposure rather than conferring high levels of resistance. These genes span diverse functional categories, including metabolism, cell envelope and cell wall processes, stress responses, and prophage associated loci, indicating that vancomycin tolerance reflects broad physiological adaptation.

Notably, the enrichment of prophage associated genes among vancomycin survival determinants, together with vancomycin induced increases in prophage associated transcripts, suggests that prophage elements may play a role in survival during antibiotic exposure.

Cross-antibiotic testing revealed that these mutants often exhibited further reduced growth under penicillin G, ciprofloxacin, and roxithromycin, indicating that the pathways involved in vancomycin tolerance also influence susceptibility to other antibiotic stresses.

Together, our findings indicate that survival under vancomycin exposure depends on the activity of multiple physiological processes rather than solely on direct modification of a drug target and underscores the complexity of bacterial stress responses that underline reduced antibiotic susceptibility.

## Acknowledgments

We would like to acknowledge Dr. Emily Goodall for further advice on transposon insertion sequencing. This work was supported by the BBSRC and the EPSRC through two grants awarded to S.P. and U.L. (BB/V008021/1, EP/Y023528/1). This work was further supported via the JPIAMR project ERADIAMR (MR/Y033892/1) awarded to S.P.. The vancomycin probe work was further supported by NHMRC (Australia) Ideas APP2004367. This project also utilized equipment funded by a Wellcome Trust Institutional Strategic Support Fund (WT097835MF), a Wellcome Trust Multi-User Equipment Award (WT101650MA) and a BBSRC award (BB/Z515942/1).

The funders had no role in study design, data collection and analysis, decision to publish, or preparation of the manuscript. The views expressed are those of the authors. For the purpose of open access, the authors have applied a ‘Creative Commons Attribution (CC BY) licence to any Author Accepted Manuscript version arising from this submission.

## Author contributions

Conceptualization, S.P.; methodology, S.Z., U.L., P.A.O.N., A.F., A.R.J., X.B., M.A.H., M.L., B.Z., M.A.T.B., A.G., and S.P.; formal analysis, S.Z., U.L., P.A.O.N., A.F., A.R.J., X.B., M.A.H.

M.L., B.Z., M.A.T.B., A.G., and S.P.; generation of figures, S.Z, and S.P.; investigation, S.Z.; resources, S.Z., U.L., P.A.O.N., A.F., A.R.J., X.B., M.A.H., M.L., B.Z., M.A.T.B., A.G., and S.P.; data curation, S.Z, and S.P.; writing – original draft, S.Z. and S.P.; writing – review & editing, S.Z., U.L., P.A.O.N., A.F., A.R.J., X.B., M.A.H., M.L., B.Z., M.A.T.B., A.G., and S.P.; visualization, S.Z., and S.P.; supervision, S.P.; project administration, S.P.; funding acquisition, U.L., and S.P..

## Data availability

The datasets generated during the study (e.g. processed datasets, analysis outputs, or supplementary source files) are included within the manuscript and supplementary information files. Further materials supporting the findings of this study are available from the corresponding author upon reasonable request. TraDIS sequencing data generated from *Staphylococcus aureus* ST398 strain S0385 transposon mutant libraries have been deposited in the NCBI Sequence Read Archive under BioProject accession PRJNA1465634. Associated BioSample accessions are SAMN59750722–SAMN59750733. Associated SRA run accessions are SRR38560739–SRR38560750.

## Methods

### Bacterial strains and growth conditions

Two *Staphylococcus aureus* genetic backgrounds were used in this study. A high-density transposon mutant library in *S. aureus* subsp. *aureus* ST398 strain S0385 was used for genome-wide TraDIS analysis. This library was generated previously, as described by Ba *et al*. (2023) (17). In addition, defined transposon insertion mutants from the *S. aureus* USA300 JE2 Nebraska Transposon Mutant Library were used for targeted phenotypic validation of selected loci. The corresponding parental strains (S0385 or USA300 JE2) were included as wild-type controls in all experiments. Unless otherwise stated, strains were grown in LB broth at 37 °C with shaking at 200 rpm as previously described (9).

### Culture conditions and DNA extraction for TraDIS analysis of S0385 library

To prepare the input pool for TraDIS analysis, 1 mL of the glycerol stock (stored at −70 °C) was added into 40 mL of LB broth supplemented with 5 μg/mL erythromycin (Em) and incubated overnight at 37 °C with shaking at 200 rpm. The following day, the culture was diluted 1:10 into 400 mL of fresh LB + 5 μg/mL Em and incubated under the same conditions until early exponential phase. The culture was then divided into three100 mL aliquots and treated with 0, 1, 10 μg/mL vancomycin. After 5 hours of incubation, cells were harvested by centrifugation at 4 °C for 20 minutes at 4,000 × g. Genomic DNA was extracted using the Qiagen Blood and Tissue Kit, including an additional lysostaphin lysis step to ensure efficient lysis of Gram-positive cells, as previously described (50).

### Sequencing library preparation and High-Throughput Sequencing

Sequencing libraries were prepared as previously described by Ba *et al*. (2023) (17) with slight modifications. Briefly, 1 μg of genomic DNA was fragmented to 200-300 bp using a Covaris E210 ultrasonicator. Fragment size distribution was validated using an Agilent 2100 Bioanalyzer with a High Sensitivity DNA kit. DNA fragments were end-repaired and ligated to Illumina-compatible adapters using the NEBNext Ultra II DNA Library Prep Kit for Illumina (New England Biolabs), following the manufacturer’s protocol. Libraries were PCR amplified using 100 ng of adapter-ligated DNA, a transposon-specific primer (17), and indexing primers from the NEBNext Multiplex Oligos for Illumina (Index Set 1) for 22 cycles. PCR products were cleaned using 0.9× SPRIselect beads (Beckman Coulter). Final libraries were pooled in equimolar ratios and sequenced as previously described (51), with some modifications. We used the p1 300 cycles flow cell on the Illumina NextSeq 1000 platform, single-end 300 bp reads, and a custom Read 1 primer (GACACTATAGAAGAGACCGGGGACTTATCAGC) for read mapping and essentiality analysis.

Sequencing reads were processed using the Bio-Tradis pipeline. Reads were aligned to the *S. aureus* S0385 reference genome (GenBank accession AM990992.1) using *bwa*, and alignments were sorted and indexed with *samtools*. Transposon insertion sites were identified using the TraDIS_gene_insert_sites script, and the number of unique insertion sites per gene was calculated for each sample.

### Comparative TraDIS analysis across conditions

For comparative analysis, the number of unique transposon insertions per gene was used as a measure of mutant representation. Genes were compared between cultures that survived exposure to vancomycin at inhibitory concentrations and those that did not. Log₂ fold changes (log_2_FC) in unique insertion counts were calculated between conditions, and statistical significance was assessed using the Bio-Tradis statistical framework. Genes were considered associated with loss of survival under vancomycin exposure if they met the following criteria: (i) |log_2_FC| > 1.5 with a significant *p* value, (ii) ≤3 unique insertions across vancomycin-surviving samples, and (iii) ≥4 unique insertions in at least one drug-free control replicate, indicating that the gene was not required for growth in LB medium.

### Gene annotation and functional analysis

Genes of interest were annotated using GenBank records and the KEGG database. KEGG orthology and pathway assignments were obtained using BlastKOALA, and conserved protein domains were identified using the KEGG SSDB motif search tool, including Pfam domain annotations, as previously described (45).

### Growth assays

For growth assays, overnight cultures were grown in LB broth at 37 °C with shaking at 200 rpm, diluted 1:100 into fresh LB, and allowed to reach early exponential phase. For cultures containing the pCM29 plasmids, the cultures were grown overnight in 10 µg/mL chloramphenicol and diluted 1:100 in fresh LB (no chloramphenicol) before the growth assays in vancomycin. Cultures were then exposed to vancomycin at various concentrations, alongside a no antibiotic control. Growth kinetics were monitored using a CLARIOstar microplate reader (BMG Labtech, Germany) by measuring optical density at 600 nm (OD₆₀₀) at regular time intervals. Plates were subjected to orbital shaking (200 rpm, 300 sec) before each reading to ensure culture homogeneity.

### Kill-Time assay

Overnight cultures were grown in LB broth at 37 °C with shaking at 200 rpm, diluted 1:100 into fresh LB, and allowed to reach early exponential phase, then diluted to approximately 10^6^ cells/mL. Vancomycin was added to the cultures at a final concentration of 1 µg/mL or 10 µg/mL. Cultures were left for five hours at 37 °C with shaking at 200 rpm and 1 mL samples were taken hourly. At each time point the samples were washed in LB to remove residual antibiotics, followed by a serial dilution and plating on LB agar plates to measure colony forming units (CFUs).

### Minimum inhibitory concentration (MIC) determination

Minimum inhibitory concentrations were determined using the broth microdilution method as previously described (48). Approximately 5 × 10⁵ cells were inoculated per well in a 96 well microplate containing LB broth supplemented with the antibiotic under investigation. Antibiotic concentrations ranged from 64 µg/mL to 0.0625 µg/mL, prepared by two-fold serial dilution. Plates were incubated at 37 °C for 18 h. Optical density at 600 nm (OD₆₀₀) was measured using a CLARIOstar microplate reader (BMG Labtech, Germany) after 18 h. The MIC was defined as the lowest concentration of vancomycin at which no visible growth was observed, as determined by comparison to positive growth control wells.

### RNA extraction and RT-qPCR

Cultures treated with vancomycin and corresponding untreated controls were harvested by centrifugation (4,000 rpm, 15 min) following 2 h of incubation with vancomycin (1µg/mL) at 37°C, 200 rpm. Cell pellets were resuspended in TE buffer (Invitrogen) supplemented with lysostaphin (Sigma- Aldrich) (final concentration 2 µg/mL) and incubated at 37 °C for 1 h. Proteinase K (2mg/mL) (Qiagen) was added for the last 15 mins of the incubation.

Total RNA was extracted using the RNeasy Midi Kit (Qiagen) according to the manufacturer’s instructions and as previously described (9), including the on-column DNase treatment step to remove residual genomic DNA. RNA concentration and purity were assessed prior to downstream processing. Two milligrams of RNA were used for reverse transcription with random hexamer primers using the High-Capacity cDNA Reverse Transcription Kit (Applied Biosystems), following the manufacturer’s protocol. No-reverse transcription (no-RT) controls were included for all samples to confirm the absence of genomic DNA contamination.

Reverse transcription reactions were carried out with an initial primer annealing step at 25 °C for 10 min, followed by cDNA synthesis at 37 °C for 120 min, and reaction termination at 85 °C for 5 min, in a final reaction volume of 20 µL. Resulting cDNA samples were diluted fivefold in 10 mM Tris-HCl (pH 8) and stored at −20 °C until quantitative PCR analysis.

Real-time qPCR reactions were performed in a total volume of 10 µL containing 1× PowerUp™ SYBR Green Master Mix (Applied Biosystems), 500 nM of each forward and reverse primer (File S7), and 5 ng of template cDNA. Amplification was carried out on a QuantStudio™ 1 Real-Time PCR System (Thermo Fisher Scientific) using the following cycling conditions: an initial uracil-DNA glycosylase (UDG) activation step at 50 °C for 2 min, followed by DNA polymerase activation at 95 °C for 2 min. PCR cycling consisted of 40 cycles of denaturation at 95 °C for 15 s and combined annealing/extension at 60 °C for 1 min.

Following amplification, melt curve analysis was performed by increasing the temperature from 60 °C to 95 °C at a rate of 0.1 °C/s with continuous fluorescence acquisition to verify amplification specificity. Threshold cycle (Ct) values were determined using Design & Analysis (DA2) software v2.6.0 (Thermo Fisher Scientific).

Transcript levels were determined by absolute quantification using standard curves generated from known gene copy numbers. Standard curves were prepared from genomic DNA extracted using the DNeasy Blood & Tissue Kit (Qiagen), and DNA concentrations (ng/µL) were measured by absorbance at 260 nm using a NanoDrop spectrophotometer (Thermo Scientific). DNA concentrations were converted to gene copy numbers per milliliter using a copy number calculator (https://www.technologynetworks.com/tn/tools/copynumbercalculator). cDNA copy numbers per mL were normalized to the corresponding *gmk* copy numbers per mL to account for differences in input material.

### Prophage induction assay (quantitative PCR of extracellular DNA)

Exponentially growing *S. aureus* S0385 cultures were incubated with either 0.5 µg/mL mitomycin C, 1 µg/mL or 0.5 µg/mL vancomycin, or left untreated as a control. Samples were collected at hourly intervals. Culture density (OD₆₀₀) was measured, after which cells were removed by filtration through a 0.2 µm pore-size filter to removes cells. Filtrates were diluted ten-fold in 10 mM Tris-HCl (pH 8.0) prior to analysis.

Quantitative PCR was performed as described previously using absolute quantification, with the modification that the initial denaturation step was extended to 10 min to break down the viral capsids.

### Measurement of vancomycin binding by flow cytometry

Vancomycin binding was assessed using fluorescently labeled vancomycin as previously described (52, 53). Briefly, a fluorescent antimicrobial derivative of vancomycin that retains vancomycin’s activity profile was added to added to cultures grown in LB at a final concentration of 2 µg/mL. At the indicated time points, 30 µL aliquots were collected, pelleted by centrifugation (12,000 rpm, 1 min), and washed once in M9 minimal salts medium ((Merck, Germany) prepared in Milli-Q water and supplemented with 2 mM MgSO_4_ and 0.1 mM CaCl_2_) to halt further antibiotic uptake (54). Cells were then resuspended in M9 medium prior to flow cytometric analysis.

Flow cytometry measurements were performed using a CytoFLEX Flow Cytometer (Beckman Coulter, USA). Cell size and granularity were assessed using forward scatter (FSC) and side scatter (SSC) channels. A medium only control was used to distinguish bacterial events from background noise. Fluorescence of individual bacteria was measured using the fluorescein isothiocyanate (FITC) channel (excitation 488 nm, emission 525/40 BP).

Instrument settings were as follows: FSC gain 1,000; SSC gain 500; Violet SSC gain 1; FITC gain 500. A threshold value of SSC-A = 10,000 was applied to limit background noise. Data acquisition was performed using CytExpert software (Beckman Coulter, USA), and data were exported and analyzed using FlowJo™ software version 10.9 (BD Biosciences, USA).

## Supplementary Figure Legends

**Figure S1.**
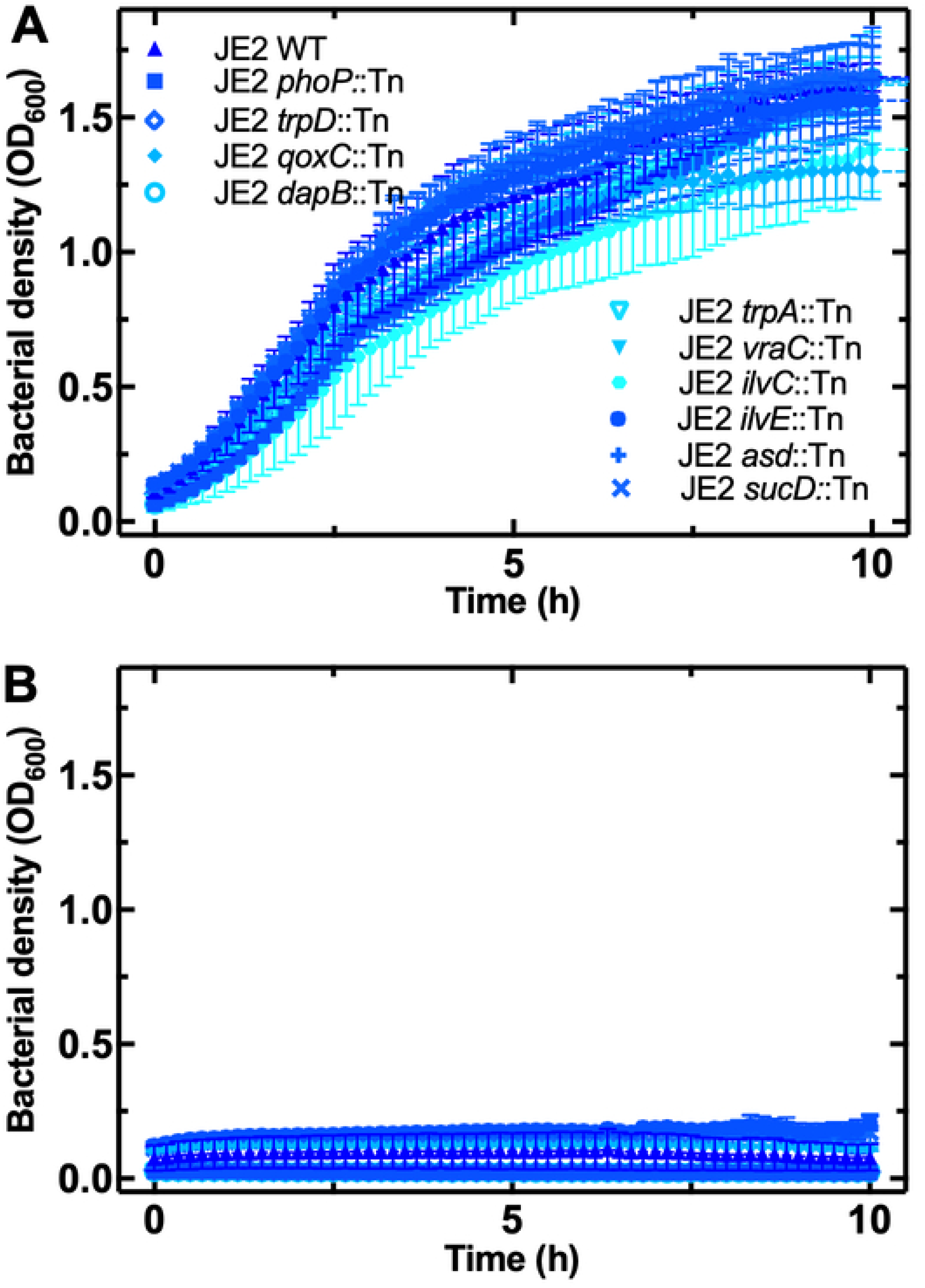
Impact of selected USA300 transposon mutants on growth during exposure to vancomycin at the MIC. Growth of defined mutants from the *S. aureus* USA300 JE2 Nebraska Transposon Mutant Library in **(A)** the absence of vancomycin and (B) the presence of vancomycin at its MIC, i.e. 2 µg/mL. Growth was monitored by OD_600_. Data represent mean and standard deviation from three biological replicates.

**Figure S2.**
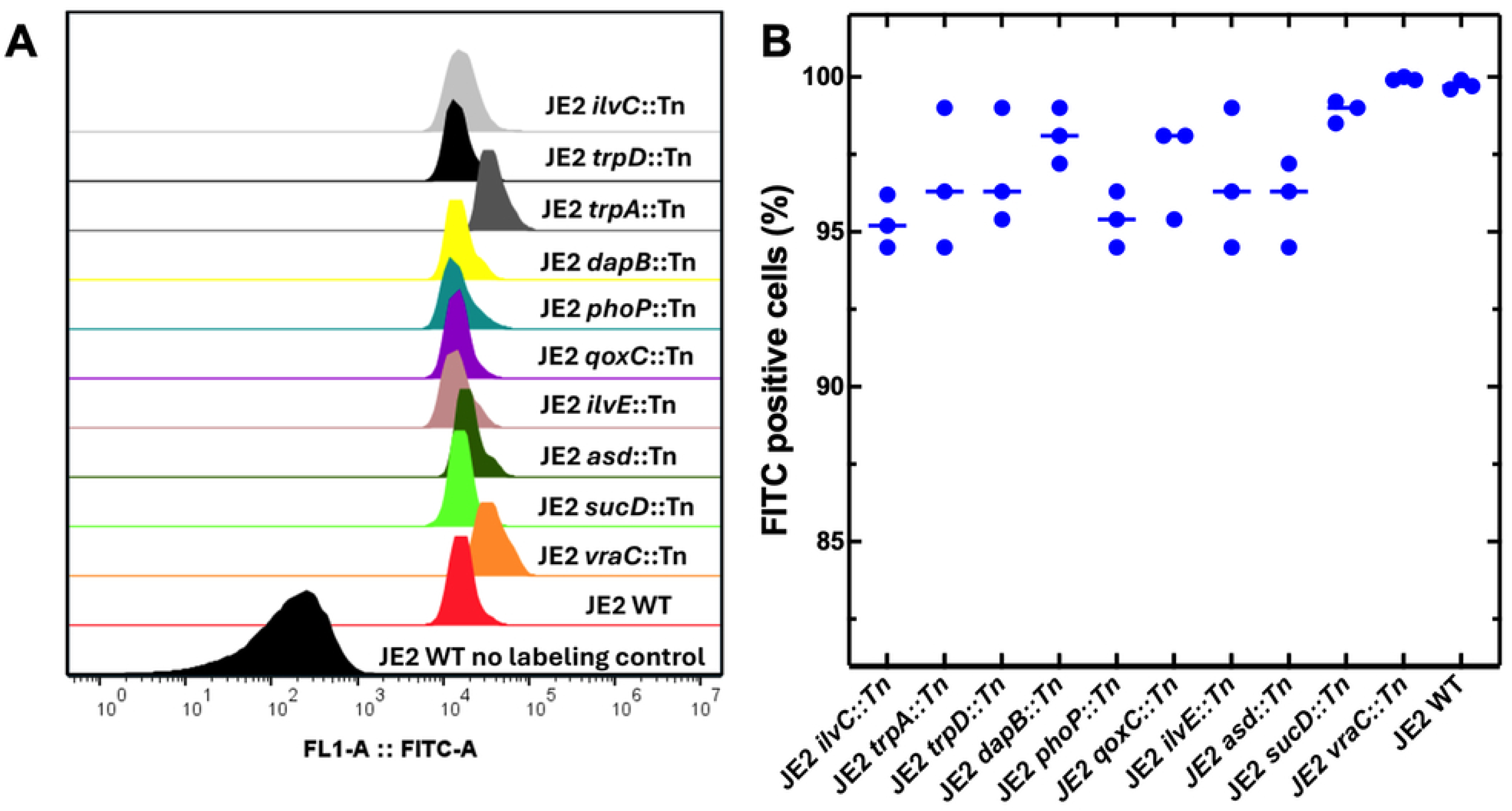
Impact of selected USA300 transposon mutants on the binding of vancomycin. **(A)** Representative histogram showing the distribution of FITC fluorescence intensity (FITC-A) for WT and transposon mutant strains following 10 min exposure to 2 µg/mL vancomycin-NBD (i.e. the MIC of vancomycin-NBD against the JE2 WT strain). Each curve represents an individual strain, illustrating vancomycin binding compared to the unlabeled WT control. **(B)** Percentage of FITC-positive cells measured by flow cytometry after 10 min incubation in 2 µg/mL FITC-labelled vancomycin. Three biological replicas shown.

## Supplementary File Legends

**File S1. Transposon insertion counts and differential analysis for cultures exposed to vancomycin at the MIC.** Genome-wide transposon insertion counts per gene, log2 fold change, and associated p values for S0385 transposon library cultures incubated for 5 h in the presence of vancomycin at the wild-type MIC.

**File S2. Transposon insertion counts for LB-grown control cultures.** Genome-wide transposon insertion counts per gene for S0385 transposon library cultures grown in LB only, derived from the same cultures prior to vancomycin exposure.

**File S3. List of genes associated with loss of survival at the vancomycin MIC.** Genes identified by TraDIS as significantly enriched in MIC-non-surviving cultures are grouped by predicted biological function based on genome annotation. Gene names are shown where available; otherwise, locus tags are provided. Genes were considered significant based on the following criteria: log₂FC > 1.5, a statistically significant p value, ≤3 transposon insertions in genes required for survival at the vancomycin MIC (File S1), and ≥4 insertions in at least one replicate under LB growth conditions, indicating that the gene is not required for growth in LB (File S2).

**File S4. Predicted Pfam domains identified in the protein sequence using KEGG annotation.** Pfam domains detected in proteins from predicted prophage regions. The table shows the prophage region, locus tag, predicted protein annotation, identified Pfam motifs, corresponding E-values, and functional notes. Pfam domain annotations were retrieved from KEGG database resources. An E-value cutoff of 0.01 was applied, and only hits corresponding to bacterial or bacteriophage proteins were considered.

**File S5. Strains from Nebraska Transposon Mutant Library (NTML) - collection generated in the *S. aureus* USA300 background used in this study.**

**File S6. Minimal inhibitory concentration (MIC) for USA300 NTML strains.**

**File S7. List of primers.** All primers were designed for *S. aureus* S0385.

